# Fos labels a distinct excitatory neuron ensemble for memory retrieval

**DOI:** 10.1101/2025.06.14.659414

**Authors:** Ning Ding, Qian Sun, Chenyun Wu, Yujie Xiao, Yiting Li, Xin Wang, Yousheng Shu, Peng Yuan

## Abstract

Experience-triggered expression of Fos has been extensively used as a marker representing memory engram. It is generally assumed that Fos-positive neurons are preferentially excited by the experience, yet this has not been directly examined. In this study, we tracked neuron activity and Fos expression in somatosensory cortex and dentate gyrus of hippocampus in mice, during electric shock-induced fear memory formation and retrieval. Contrary to the prevailing view, we found that calcium features fail to predict Fos for individual cells. Fos-positive neurons show no specific activation to stimuli during memory encoding, but their activities become stimuli-selective during retrieval. Surprisingly, inhibiting neurons that will express Fos during encoding does not affect the induction of memory-guided behavior upon their activation. Network simulation and voltage imaging further showed that connectivity to interneurons, not stimulus responsiveness, is a major factor determining Fos expression. Our results indicate that Fos labels excitatory neurons specific for memory retrieval.

## Main

Memory is a fundamental cognitive process that shapes behavior based on prior experiences. A key concept in memory formation involves the emergence of engram, which was first introduced over a century ago by Richard Semon^1^. Memory engram refers to the physical and functional changes in the brain that happen during the encoding of memory^2^. Over the past few decades, neurons expressing immediate early genes (IEGs) have been identified as a physical substrate of memory engram. This was prompted by the pioneering work of Reijmers et al. that described the activation of IEG promoter during memory encoding^3^. Subsequent work by Liu et al. tagged IEG (*Fos*)-expressing neurons in the dentate gyrus of hippocampus, specifically during memory encoding with optogenetic constructs and demonstrated that activation of these tagged neurons led to behavioral expression of fear memory retrieval^4^. Since then, similar observations have been made in numerous laboratories^5,6^. *Fos*-based labeling is now used as a proxy to identify engram in memory research. However, the relationship between neuron activity patterns and Fos expression remains poorly understood.

The prevailing hypothesis regarding why some neurons express Fos during memory encoding posits that they are highly activated by the experience to be memorized, thereby providing an “information tag” for the memory. This view is supported by the molecular pathway that regulates *Fos* transcription. Neuron activity causes elevated intracellular calcium (Ca^2+^), leading to activation of calmodulin-dependent protein kinase^7^ and phosphorylation of cAMP-response element binding protein (CREB) in order to induce *Fos* expression^8^. In addition, neurons exhibiting elevated excitability show high levels of CREB^8^, suggesting that they are primed for Fos expression upon activation during memory encoding. This hypothesis also suggests that reactivation of the Fos-labeled neurons could reinstate the “information tag”, thus serving as a neural mechanism for memory retrieval.

Based on the above hypothesis, it has been widely assumed that experience-related neuron activity triggers the induction of Fos expression and thus links Fos-expressing cells to specific memories, yet this has not been directly tested. This is due to the technical challenges of performing *in vivo* imaging of neuron activities coupled with cognitive tasks, and stably tracking the expression of Fos at single-cell resolution. Several recent studies identified a wide range of heterogeneity between neuron activity and Fos expression^9^. Yet these studies did not directly examine activities during memory encoding, thus the relationship between information representation and Fos expression remains untested. To address this gap of knowledge, we conducted *in vivo* imaging of neuron activities and Fos expression during fear memory encoding and retrieval in somatosensory cortex and dentate gyrus of hippocampus. Contrary to the conventional assumptions, our data revealed that Fos-expressing neurons do not show experience-related activity during encoding, but their representation became memory-related during subsequent retrieval. Intriguingly, we further discovered that activities of the Fos-expressing neurons are not essential during encoding for linking them with specific memories. Our finding strongly indicate that memory encoding and retrieval may engage different neural populations, with Fos preferentially labeling retrieval-related populations (memory ecphory). These results have implications for dissecting the neural mechanism underlying memory formation.

## Results

### Overall heightened calcium activities exist in neurons prior to Fos expression

The conventional view hypothesizes that Fos induction is induced by elevated intracellular calcium (**Figure 1a**, process 1) due to high neuron activities (**Figure 1a**, process 2) elicited by the stimulus or experience to be memorized (**Figure 1a**, process 3). In this study we systematically examined all these processes, and we first studied the relationship between neuronal calcium transients and Fos expression. We employed an imaging approach to simultaneously monitor these two factors, by injecting adeno-associated virus (AAV) for GCaMP6f expression in mice crossbred from TRAP2 (Fos^2A^-iCreER) and Ai9 strains (**Figure 1b-c**). To identify neuron activities corresponding to Fos-driven tdTomato expression a few days later, we aligned the observed neurons across different sessions (**Figure 1c**). For inducing event-associated Fos expression, we applied a unilateral electric shock to the hindlimb of the mice, which caused elevated Fos expression in the contralateral somatosensory cortex compared to the ipsilateral side (**Extended data figure 1**). With this setup, we imaged calcium activities in somatosensory cortex neurons during electric shock, and compared calcium transient features in neurons with and without Fos expression (**Figure 1e-f**). We examined transient width, amplitude, frequency, and cumulative calcium levels, and found that Fos-positive neurons exhibited significantly increased transient width compared to Fos-low neurons, with spike counts and total calcium influx also showed increased trends (**Figure 1f**). In addition, we imaged activities and Fos expression in dentate gyrus neurons using a dual-color miniaturized microscope (miniscope) in freely moving mice during application of electric shocks. Again, we observed a trend of widened calcium activities in Fos-positive neurons (**Figure 1g-h**). Consistent with previous reports^10^, these data indicated that neurons exhibited overall elevated calcium activities prior to Fos expression.

**Figure 1.**
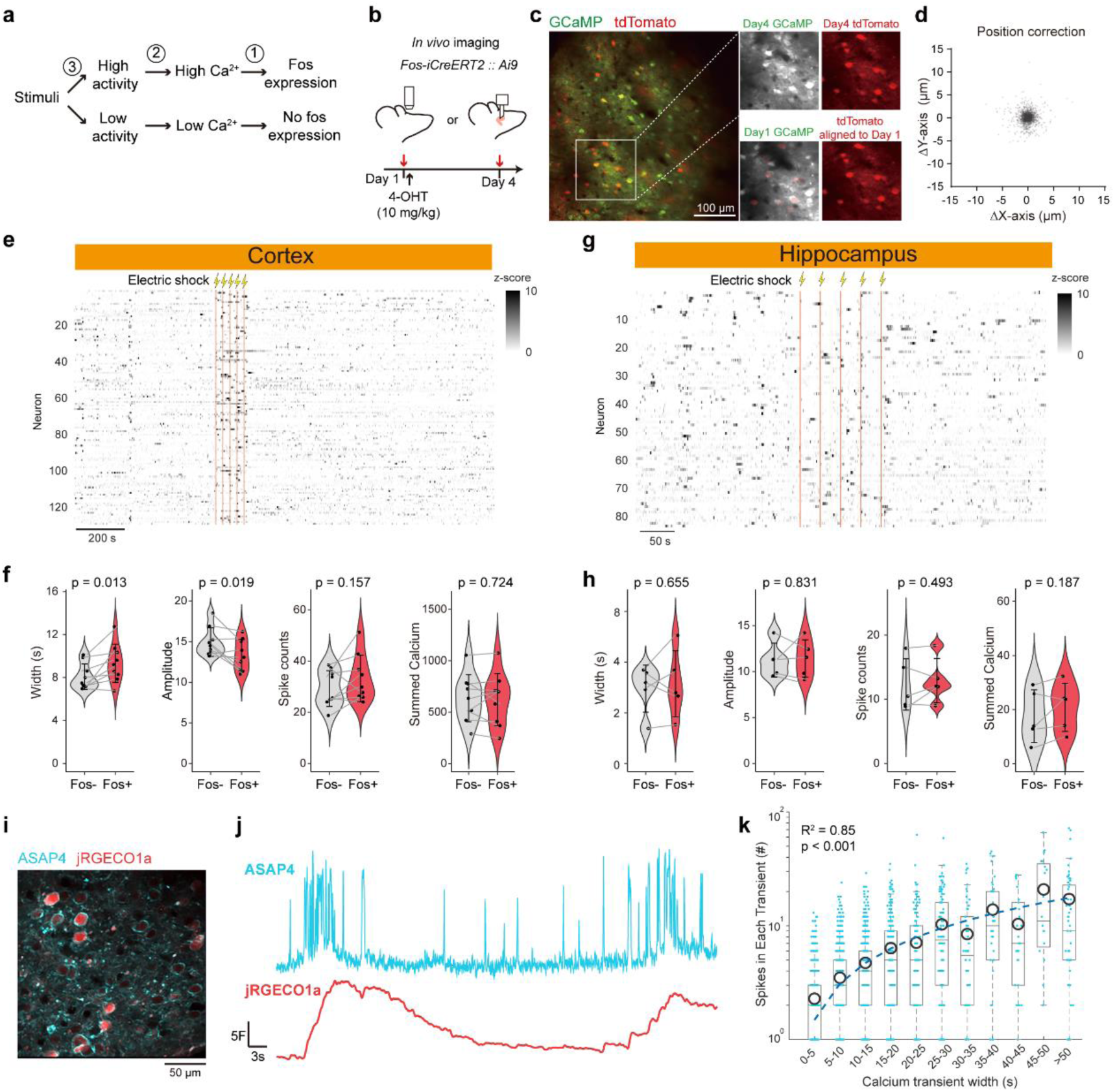
Overall increase in calcium activity in fos-positive neurons. **a**, Conventional hypothesis of Fos expression. Numbers indicate three different aspects on this causal link are tested in the current study. **b**, Schematics of two-photon calcium imaging experiments. **c**, Representative two-photon microscopy images showing alignment of neurons within the field of view across sessions. Right panels show zoomed-in views from the box on the left. **d**, The scatter plot depicting alignment errors between neurons across two sessions. **e,** Representative calcium responses to the electric shock from neurons in the somatosensory cortex. Orange bars indicate time periods of electric shocks. **f,** Quantification of calcium activity parameters of fos-positive and fos-negative neurons, including width, amplitude, spike counts, and summed calcium. Two-sided paired-sample t test, n = 9 mice. **g**, Representative calcium responses to the electric shock from neurons in the DG. Orange bars indicate time periods of electric shocks. **h**, Quantification of calcium activity parameters similar to **f**. Two-sided paired-sample t test, n = 5 mice. **i-j**, Representative two-photon images (**i**) and traces (**j**) of simultaneous recording of membrane voltage and intracellular calcium. **k**, Quantification of the relationship between calcium transient width and spike counts during each transient. n = 5513 events. Pearson’s correlation was used for statistics.

In order to investigate whether elevated calcium activities associated with Fos expression reflect enhanced neuron firing or changes in calcium-related signaling (related to process 2 in **figure 1a**), we performed simultaneous imaging of membrane voltage and intracellular calcium *in vivo*. We employed a voltage sensor ASAP4b with relatively slow kinetics^11^ and combined with a calcium sensor of red fluorescent emission (**Figure 1i**). This combination allowed us to record voltage spikes from 50-100 neurons at a frame rate close to 100 Hz, achievable with commercial two-photon microscopes. We were able to identify time periods of neuron firing that were difficult to resolve based on calcium traces (**Figure 1j**). By grouping activity events based on different calcium transient widths, we found a strong linear correlation between the number of spikes and calcium transient widths (**Figure 1k**), indicating that elevated calcium activities associated with Fos expression are likely resulted from enhanced neuron firing in these cells. Together, these data indicate that Fos expression is related to elevated neuron activities.

### Experience-induced calcium activities poorly predict Fos expression in individual neurons

The above analysis indicated a correlation between heightened neural activities and Fos expression. To quantitatively evaluate the robustness of this correlation at individual cell level, we constructed support vector machine (SVM) classifiers for Fos expression using calcium transient parameters (**Figure 2a**). We explored all possible combinations of several calcium transient parameters and found that, although some models showed performance statistically superior than those from shuffled control, even the full model incorporating all parameters yielded an accuracy of less than 60% (**Figure 2b**). Training a convolutional neural network based on time sequences of calcium traces did not further improve classification performance (**Figure 2c**). These findings suggest that increased calcium activity is a weak predictor of Fos expression at single-cell level. To further visualize this weak correlation, we performed dimensionality reduction on the calcium transient parameters using UMAP (Uniform Manifold Approximation and Projection)^12^. We pseudocolored individual neurons on the UMAP based on the predicted Fos expression probability, and marked cells of experimentally verified Fos expression (**Figure 2d, e**).We observed that vectors from the centroid of Fos-negative neurons to Fos-positive neurons aligned with the direction of predicted Fos expression (**Figure 2f**). The probability of predicted Fos expression closely correlated with experimental observation in large groups of neurons, yet the correlation disappeared on individual cell level (**Figure 2g**). Together, these single-cell analysis showed a high degree of variance between calcium activity and Fos expression, providing evidence against a simple causal relationship between heightened neuron activity and Fos expression (processes 1 and 2 in **figure 1a**).

**Figure 2.**
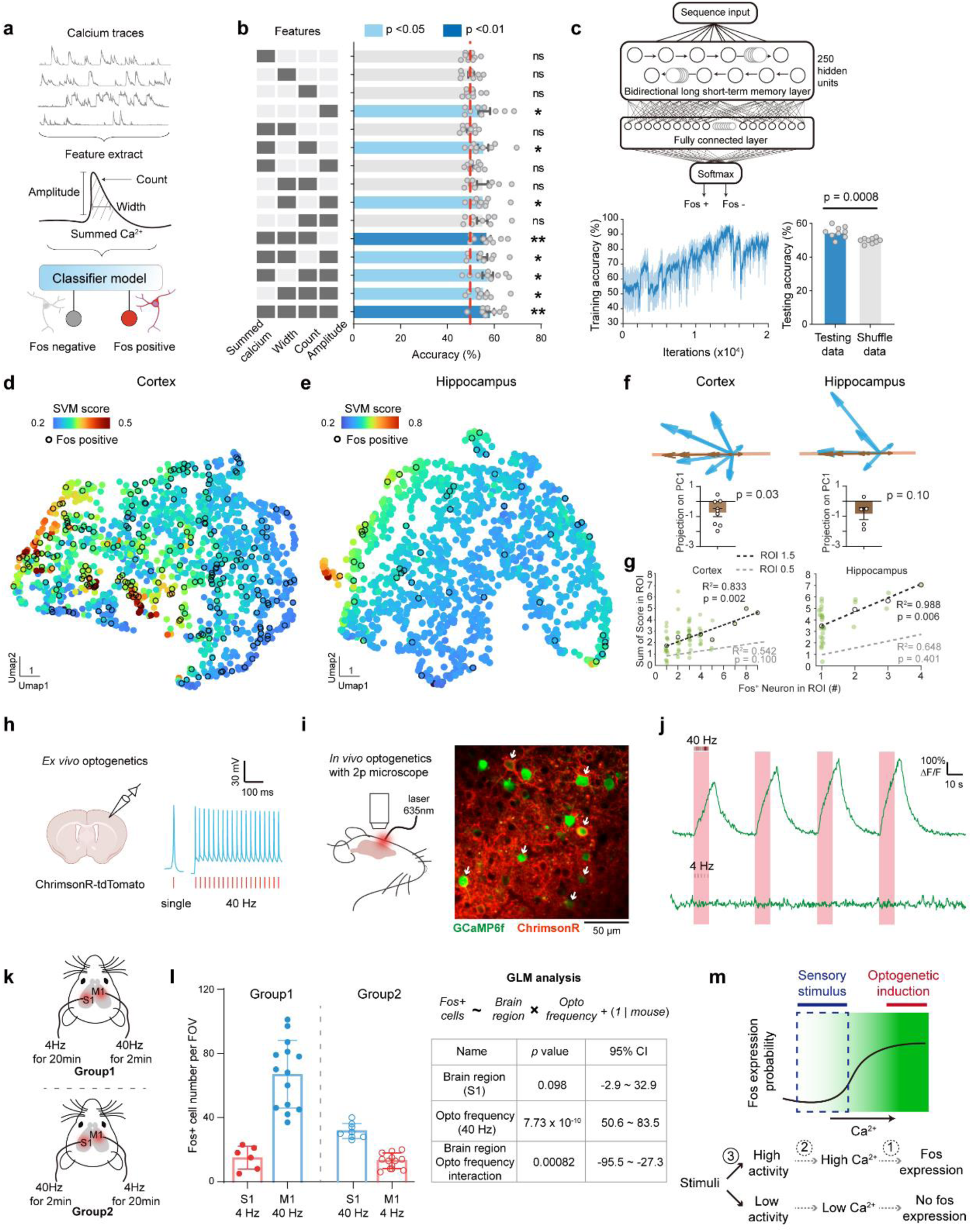
Physiological calcium rise does not predict Fos expression on single-cell level. **a**, Diagram of classifier modeling using calcium transient features. **b**, Model performance for predicting Fos expression with different inputs. One-way ANOVA test with post-hoc analysis and Dunnett’s correction for multiple comparison. *: P <0.05, **: P <0.01, n.s., not significant. n = 9 mice. **c**, Neural network classifier for Fos expression using calcium time sequences. Lower left panel shows representative training performance of the network. Dark line indicates average accuracy from a 50-iteration moving window. Lower right panel shows quantification of the performance on testing dataset. Two-side t test, p = 0.0008. n = 9 mice. **d-e**, Visualization of calcium transient features of individual neurons from cortex (**d**) and DG (**e**) in a reduced dimension, with each neuron color-coded with the predicted probability of Fos expression. Black circles indicate experimentally observed Fos expression. **f**, Vectors from the centroids of fos-negative to fos-positive cells on the UMAP. One sample t-test compared to 0, n = 9 mice for cortex and 5 mice for hippocampus. **g**, Scatter plots showing predicted and experimentally measured probability of fos expression, within sliding windows of 1.5 units on the UMAP. Dashed lines indicate linear regression with Pearson’s correlation analysis. Gray dashed lines show the same analysis for the sliding window of 0.5 units. **h**, Representative patch-clamp recording of ChrimsonR-expressing cells in acute brain slice shows faithful induction of action potentials (blue curves) with light pulses (red bars). **i**, Representative field of view for simultaneous optogenetics stimulation and in vivo two-photon imaging. Arrows indicate cells with co-expression of both constructs. **j**, Representative calcium traces (green) with 40Hz or 4Hz laser pulse stimulation. Red shades indicate time periods with stimulation laser on. **k**, Schematics of the optogenetic stimulation experiments design. **l**, Quantification of Fos expression cells in different brain regions after optogenetic activation (40Hz for 60 seconds, 4Hz for 600 seconds). Each dot indicates one brain slice. Data are represented as mean ± S.D. The analysis with generalized linear model is presented on the right panel. F-test was used to compare modeled coefficients to 0. **m**, Diagram describing the relationship between Fos expression and intracellular calcium flux. Lower diagram summarizes the poor correlation (dashed arrows) between neuron activity, intracellular calcium concentration, and fos expression.

As numerous previous *in vitro* studies have established that heightened intracellular calcium was capable of inducing Fos expression^7^, the poor correlation we observed above was unexpected. Thus, we further tested the sufficiency of high calcium levels in inducing Fos expression *in vivo*. To this end, we utilized an optogenetic approach and established that laser pulse trains induced robust calcium transients, while single pulse stimulation elicited sparse action potentials without observable calcium events (**Figure 2h-j**). We compared the efficacy of Fos induction using these two protocols, and the results showed that optogenetically induced calcium transients significantly upregulated Fos expression, while the same total number of sparsely distributed light pulses, which did not induce calcium transients, failed to produce this effect (**Figure 2k-l, Extended data figure 2**). These data indicate that extremely high intracellular calcium levels can be a powerful driver for Fos expression in neurons. However, experience-induced neuronal activity only weakly elevates calcium levels in somatosensory cortex and DG of hippocampus, thus it is not a strong determinant for Fos expression (**Figure 2m**), suggesting an even weaker correlation between experience representation and Fos expression.

### Fos labeling in the cortex does not reflect stimulus coding during initial experience

We next directly examined whether representation of external stimuli was related to Fos expression (process 3 in **figure 1a**). Using the paradigm of hindlimb electric shock we described above, we observed that some neurons in the contralateral somatosensory cortex exhibited highly selective calcium responses immediately following the application of electric shock (**Figure 3a-b**). However, upon cross-examining the expression of Fos-induced tdTomato fluorescent protein, we found no correlation between fluorescent intensity and magnitude of calcium responses to electric shock (**Figure 3a**). To formally quantify the relationship with shock responsiveness, we calculated a responsiveness index for each neuron, taking into account both response amplitude and timing (see Methods for detail). When we plotted the responsiveness against Fos-induced tdTomato fluorescent intensity, we did not find a positive correlation, and there was no difference of responsiveness in Fos-positive and Fos-negative neurons (**Figure 3c**). And neuron groups with high or low responsiveness indices did not differ in Fos expression probability (**Figures 3d**). Consistently, the distribution of mutual information between electric shock and neuron activation overlapped between Fos-positive and Fos-negative neurons (**Figure 3e**). These data strongly indicate that there is no correlation between stimulus coding and Fos expression in somatosensory cortex.

**Figure 3.**
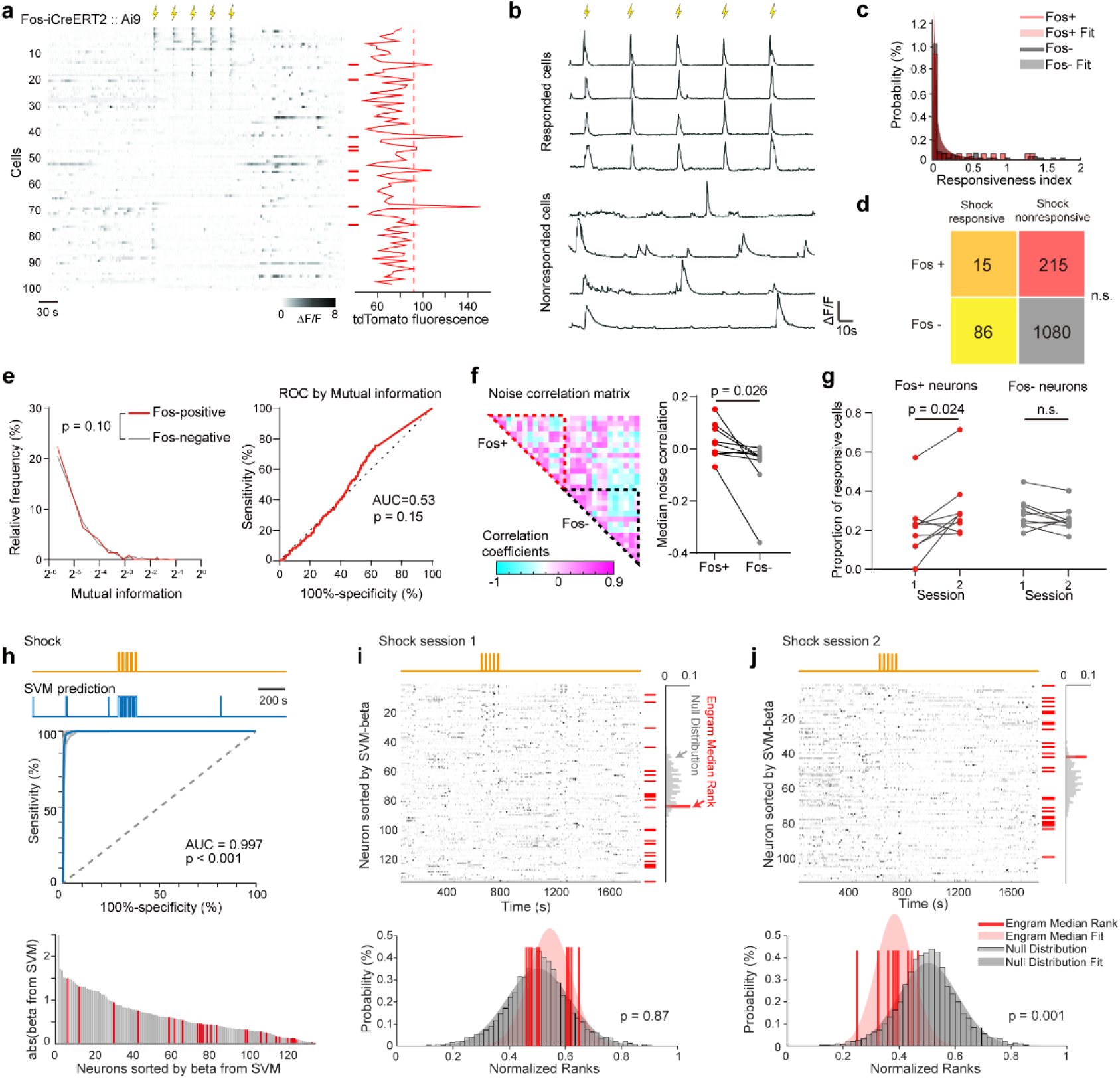
Fos-positive neurons do not encode stimulus initially but gain responsiveness later. **a**, Raster plots of raw calcium traces from a subset of imaged neurons. Cells were ranked by their responsiveness to the electric shock. Yellow flash signs indicate the time of electric shock. Right plot shows the fluorescence intensity of fos-driven tdTomato of the corresponding cells. Fos-positive cells (intensity above threshold indicated by the red dash line) are labeled with red bars. **b**, Representative calcium traces from shock responsive and non-responsive cells. Yellow flash signs indicate the time of electric shock. **c**, Histogram of the responsiveness index in fos-positive and fos-negative neurons. Shades indicate fitting results of log-normal distributions. **d**, Number of Fos-positive and Fos-negative neurons that responded and did not respond to electric shocks. Chi-square test: not significant. **e**, Left panel, the distribution of mutual information between cell activity and electrical shocks in Fos-positive and Fos-negative neurons. Kolmogorov-Smirnov test, not significant. Right panel, ROC curve for mutual information between Fos-positive and Fos-negative neurons. **f**, Quantification of the noise correlations among Fos-positive and Fos-negative neurons. Two-sided paired-sample t test, n = 9 mice. **g**, Quantification of the proportion of cells responding to electrical shocks across two sessions among Fos-positive and Fos-negative cells. Two-sided paired-sample t test, n = 9 mice for each group. **h**, Representative traces of actual periods of electric shocks and predictions from a SVM model. Lower graph shows the SVM beta coefficients ranked from high to low, with fos-positive neurons colored in red. **i**, Example of neuronal activities from the first imaging session, ranked by the neurons’ SVM coefficients, with fos-positive neurons labeled with red bars. Right panel shows the medium rank from fos-positive neurons, and null distribution generated by random sampling the same number of neurons as fos-positive neurons. Lower panel shows the quantification of medium rank from individual mice, with shades indicate fitting results of normal distribution. Rank-sum analysis, n = 9 mice. **j**, The same analysis as in **i**, using data from the second imaging session.

As our data suggest that Fos expression is not determined by stimulus-related response, we next tested whether Fos-expression is determined by the neuron’s intrinsic excitability. We measured functional connectivity (measured via noise correlation) among imaged neurons, and found that Fos-positive neurons showed higher inter-connectivity compared to Fos-negative neurons (**Figure 3f**). In addition, we analyzed the time periods before electric shock in the same dataset, and found that Fos-positive neurons showed widened Ca2+ transient durations during that time window as well (**Extended data figure 3**). And in dataset acquired without electric shock, we also found that Fos-positive neurons showed increased cumulated total calcium (**Extended data figure 3**). These data indicate that the enhanced calcium responses prior to Fos expression were not related to stimulus, but were likely due to elevated intrinsic excitability.

Several previous studies have reported selective synaptic modifications following Fos expression ^13^, thus we further examined the responsiveness of the same neuron population to electric shock applied at the same condition several days after the initial shock. Interestingly, we found that Fos-positive neurons became more sensitive to the shock during the second imaging session, despite not being preferentially responsive initially (**Figures 3g**). The increase in the proportion of shock responding neurons was only observed among Fos-positive neurons, whereas Fos-negative neurons did not show this change (**Figure 3g**). To formally quantify this shift in stimulus coding, we constructed linear SVM models to predict shock periods based on normalized calcium activities. The SVM model showed a high level of accuracy, indicating that the linear combination of individual neuron’s activity faithfully represented information related to shock (**Figure 3h**). To assess their importance in stimulus coding, we calculated the coefficients of Fos positive neurons, and compared them against a null-hypothesis distribution generated from random sample of the same number of neurons from the same dataset. This analysis revealed no difference using data acquired from the session when mice initially received electric shock (**Figure 3i**). In contrast, when applying the same process on the same group of neurons for the second electric shock session, we found that Fos-positive neurons became stimulus specific, as they showed significantly higher ranks in the classifier coefficients compared to random samples (**Figure 3j**). Overall, these results suggest that while Fos-expressing neurons do not show stimulus specificity initially, they, likely through undergoing plasticity change, become more selective in their response to stimuli over time.

### Fos-labeled DG neurons are not involved in memory encoding but are recruited for retrieval

We next sought to directly examine the relationship between neuron activity and Fos expression within the context of learning and memory, specifically in a brain region that previously used Fos labeling to represent memory engram. We performed live imaging of neuron activity and Fos expression in the DG of hippocampus during contextual fear conditioning (**Figure 4a**). We utilized a dual-color miniscope that enabled simultaneously visualization of calcium activity and Fos-driven tdTomato expression (**Figure 4b**). In order to image DG, we aspirated the cortical tissue above^14^. And we confirmed that mice with surgery were still capable of learning contextual-specific fear memory (**Figure 4c**).

**Figure 4.**
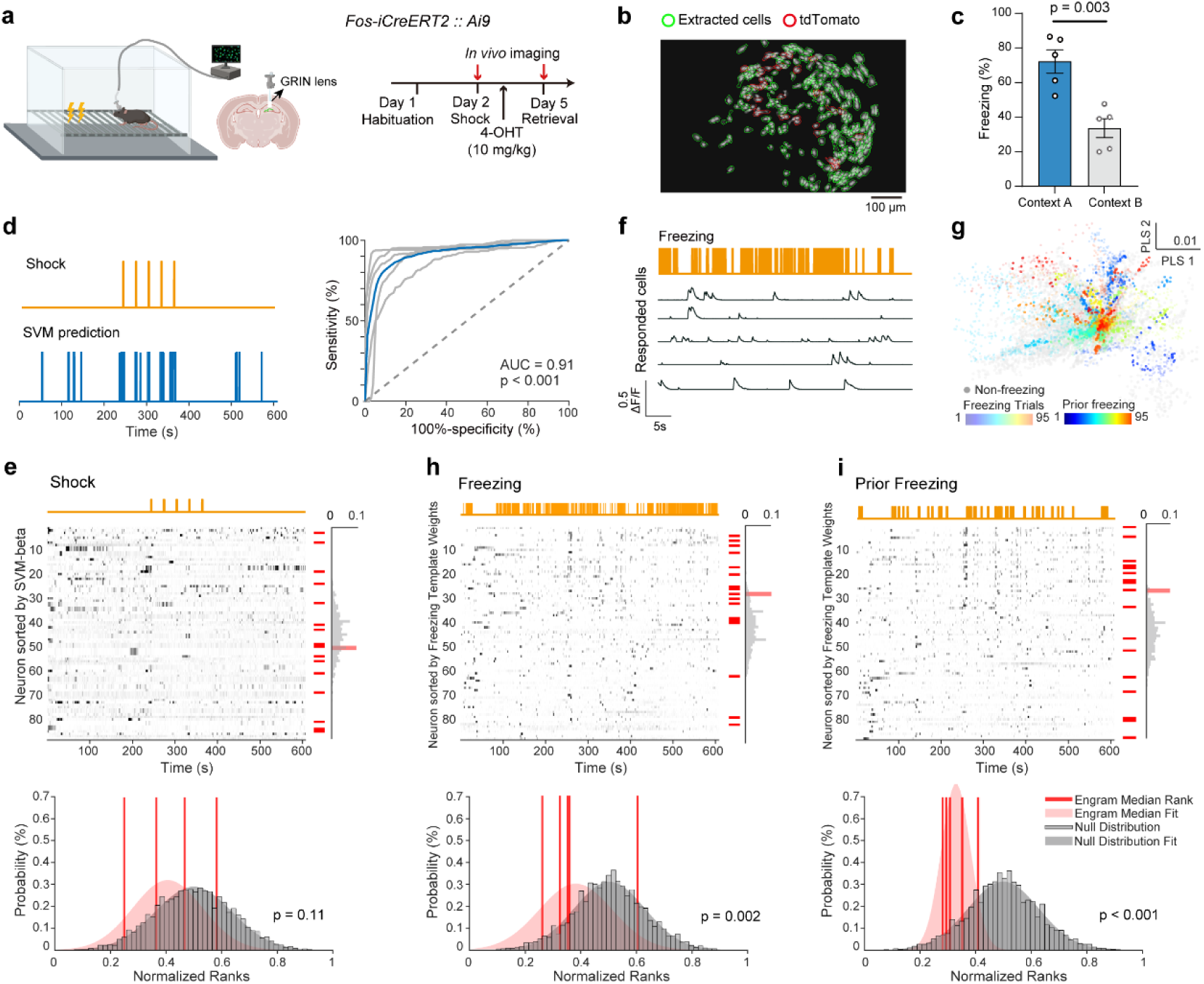
Fos-positive neurons represent memory retrieval. **a**, Schematics and timeline of experiment design. **b**, Representative image of the field of view with GCaMP and fos-driven tdTomato expression. **c**, Quantification of mice freezing behavior in environments A and B. Data are represented as mean±S.E.M. Two-sided unpaired t test, n = 5 mice. **d**, Representative traces of actual periods of electric shocks and predictions from a SVM model. Right panel shows the ROC curves for SVM prediction scores and electric shocks. The prediction of electric shock trace by neuron activity with SVM model. **e**, Example of neuronal activities from the conditioning session, ranked by the neurons’ SVM coefficients, with fos-positive neurons labeled with red bars. Right panel shows the medium rank from fos-positive neurons, and null distribution generated by random sampling the same number of neurons as fos-positive neurons. Lower panel shows the quantification of medium rank from individual mice, with shades indicate fitting results of normal distribution. Rank-sum analysis, n = 5 mice. **f**, Example traces of neuronal activities (black lines) aligned with freezing episodes (orange line). **g**, Scatter plots of the testing session neuron activity in reduced dimensions constructed using partial least square regression towards freezing. Each dot represents a timepoint and is color coded to show the temporal order of the freezing episodes (transparent) and the prior freezing periods (solid). **h-i**, Similar analysis as in **e**, using neuron activities in the testing session for time periods during freezing (**h**) or prior to freezing (**i**). Average activity during the corresponding periods is used as templates. Rank-sum test, n = 5 mice.

To investigate the neuron activity associated with memory encoding, we analyzed data acquired on the conditioning day, focusing on the activity associated with electric shocks. While Fos-positive neurons showed a general trend of elevated activity (**Figure 1**), we did not find shock responsive neurons as those in somatosensory cortex (**Extended data figure 4a-b**). To search for shock-related information in neuron population, we trained an SVM model to predict shock occurrences using recorded neuron activities. The model predicted the shock periods with high accuracy (**Figure 4d**), suggesting that the population activity in DG conveyed the information related to shock. Using analysis of SVM coefficient ranks similar to that applied in cortical neurons, we found that the coefficient ranks of Fos-positive neurons were not different from randomly sampled subsets from the same dataset (**Figure 4e**), indicating a lack of selective activation in Fos-positive neurons during memory encoding. We have also examined the neuron activities during the habituation day and found that Fos-positive neurons did not show heightened activity in the context to be paired with electric shocks (**Extended data figure 4c**). These data indicate a lack of selective activation in cells that subsequently express Fos during memory encoding.

In exploring signals associated with memory retrieval, we examined the calcium activities during the test session (**Figure 4g**). While no feature indicates precise periods of retrieval, we hypothesized that memory retrieval might be more prevalent during the animal’s freezing behavior. In addition, when we visualized the neuron activity in reduced dimensions, we noticed that time periods immediately prior to the onset of freezing were clustered (**Figure 4g**), indicating a stereotypical activity pattern that could potentially signal memory retrieval. We then used the average neural activity during these two time windows (during freezing and prior to freezing) as potential neural signature for memory retrieval, and used the levels of activity from each cell in these periods as their weights during this process. Applying the same rank analysis, we found that Fos-positive neurons showed significantly higher weights compared to randomly sampled cells in both freezing and freezing-prior periods (**Figure 4h-i**). These results indicate that Fos-positive neurons are preferentially recruited during periods likely associated with memory retrieval. In support of this view, optogenetic activation of Fos-positive neurons in a neutral context induced freezing behavior (**Extended data figure 5**), consistent with previous findings^4^. Furthermore, while we did observe a small fraction of cells responding to both electric shock and freezing, the probability of Fos expression among these cells was disproportionally low (**Extended data figure 4e**), suggesting that the coding consistency did not determine the Fos expression.

### Activity during memory encoding is not necessary for Fos-positive ensemble assignment

The above data indicate that experience-triggered calcium activity does not adequately account for Fos expression in individual neurons, and memory-encoding neurons might be a distinct population from the Fos-expressing neurons. However, it is still possible that unique patterns of activities are triggered by the stimuli that determine the expression of Fos, which cannot be resolved by calcium imaging. Thus, we next tested whether suppressing neuron activities during memory encoding blocks the recruitment of these neurons into the memory-related Fos ensemble. To achieve this, we leveraged previously reported findings that optogenetically activating a random subset of neurons prior to fear conditioning could assign them as a Fos-expressing ensemble to represent fear memory^10^. This strategy allowed us to gain genetic access to the Fos-positive ensemble during memory encoding.

We used an AAV construct that co-expressed inhibitory GtACR2 (activated by blue-light) and excitatory ChrimsonR (activated by red-light)^15^ (**Figure 5a-b**). Electrophysiological recordings in acute brain slices confirmed that bidirectional activity control could be achieved in the infected neurons with blue or red light stimulation (**Figure 5c**). We injected this virus bilaterally into the DG of hippocampus (**Extended data figure 6**), and applied optogenetic activation (30 s at 20 Hz) 5 minutes before fear conditioning (**Figure 5d**). When we reactivated these cells in a novel context the following day, we were able to induce freezing behavior, consistent with previous reports^10^ (**Figure 5e**). The induced freezing behavior closely matched the timing of light stimulation, and the effect was absent in a control group when optogenetics-induced Fos expression occurred 24 hours prior to fear conditioning (**Figure 5d-e**), indicating that this procedure successfully assigned fear memory-related information to an artificially defined neuron ensemble. We then performed the same optogenetic induction of Fos-positive ensemble 5 minutes before fear conditioning, and applied optogenetic inhibition of the same group in time periods of electric shocks (**Figure 5d**). Remarkably, we found that the inhibition did not affect the induction of freezing behavior by these neurons on the next day (**Figure 5e**). Optogenetic activation of these cells in the following day led to rapid transitions to freezing behavior, but did not affect the peak speed during periods with locomotion (**Figure 5f**), indicating a specific induction of memory-guided freezing. And the freezing levels within 20 seconds before and after turning on the laser were comparable to those in the group without optogenetic inhibition during conditioning (**Figure 5g**). These data indicate that neuron activities during memory encoding are not necessary for the assignment into the Fos-positive ensemble that represents the memory (**Figure 5h**).

**Figure 5.**
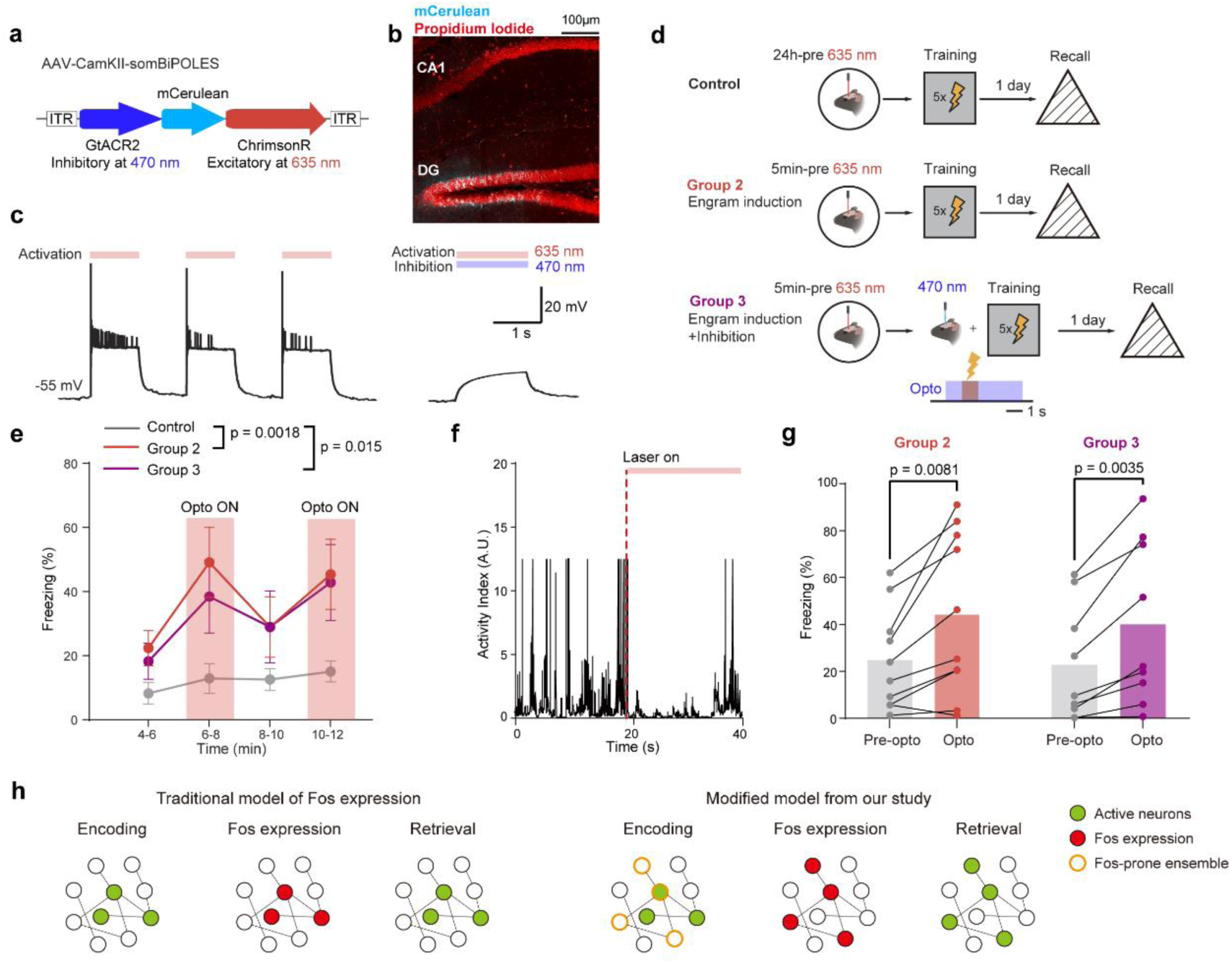
Silencing Fos-positive neurons during memory encoding does not affect linkage with fear memory. **a**, Schematics of the virus design for bidirectional optogenetic control. **b**, Representative confocal image of the virus infection in the hippocampal DG region. White dash line indicates the location of optic fiber. **c**, Representative traces of patch-clamp recordings from neurons infected with the bidirectional optogenetic construct, with red and blue laser stimulation. **d**, Schematics of the experimental design and timelines. **e**, Quantification of the freezing behaviors in different groups with (red shades) and without optogenetic activation. Data are represented as mean±S.E.M. Two-way ANOVA is used for statistical comparison, with Tukey’s multiple comparison test for post-hoc analysis. n = 6, 10, 9 mice for group 1, 2, and 3, respectively. **f**, Representative locomotion traces before and after turning on the optogenetic laser (red dashed line). **g**, Quantification of the freezing time in the test session with and without optogenetic activation. Paired t test, n = 10 and 9 mice for group 2, and 3. **h**, Diagram of the relationships among neuron activity during memory encoding, fos expression, and activity during memory retrieval.

### Inhibitory neurons may link stimulus with Fos-positive ensemble

If neuron activity during memory encoding is not necessary for the formation of Fos-positive ensemble, how would the neurons establish specific link with the event? To probe this question, we simulated a simple network with the classic Leak-Integrate-Fire model ^16^, with 30 inhibitory neurons and 70 excitatory neurons (**Figure 6a**). In order to model electric stimulus, we provided five short input current to 20 randomly selected neurons, and we further simulated intracellular calcium fluctuations with a slow decay kinetics (**Figure 6b**, see methods for details). Similar to our observation with in vivo imaging (**Figure 3**), we found several neurons that responded to the stimulus yet the calcium rise did not reach the threshold for Fos expression. On the other hand, due to high firing frequency in the inhibitory neurons at baseline, several inhibitory neurons that responded to the stimulus generated high calcium responses and became Fos positive. Fos expression then caused increased excitability and strengthened synaptic connections in these cells. Upon an offline broad activation (input current to 80 neurons, similar to broad activation during sleep^17^), we observed differential activity profiles among excitatory neurons with some generated above-threshold calcium. This model reproduced our experimental observation that Fos-positive excitatory ensemble is a distinct population from the stimulus responsive neurons (**Figure 6c**). Furthermore, this model generated two specific predictions: 1) Stimulus responsiveness should show good correspondence with Fos-expression in inhibitory neurons; and 2) Synaptic connection between Fos-positive inhibitory ensemble and Fos-positive excitatory ensemble should be relatively weak (**Figure 6d**).

**Figure 6.**
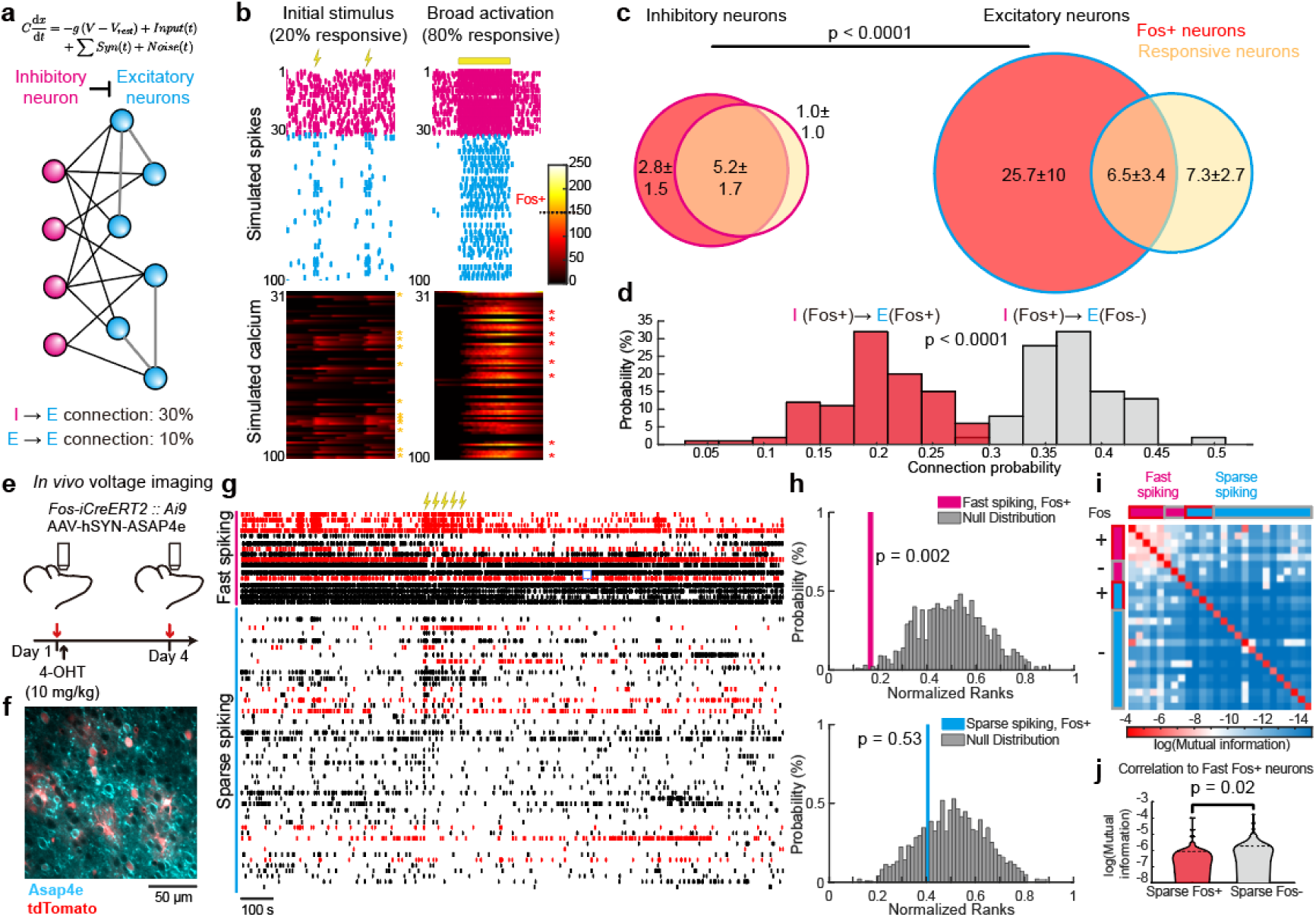
Silencing Fos-positive neurons during memory encoding does not affect linkage with fear memory. **a**, Schematics of neuronal network simulation. **b**, Representative simulated spike trains and calcium activities during the initial stimulation (yellow flash signs) and off-line activation (yellow bar). Asterisks indicate stimulus-responsive cells (yellow) and Fos-positive cells (red). **c**, Venn diagrams of Fos-positive and stimulus-responsive neurons in simulated inhibitory and excitatory groups. Data are represented by mean ± standard deviation with 100 simulations. Chi-square was used for statistical comparison. **d**, Distributions of connectivity to inhibitory Fos-positive neurons. Two-tailed t-test was used for statistical comparison. n = 100 simulations. **e**, Timeline of the *in vivo* voltage imaging experiment. **f**, Representative field of view for two-photon voltage imaging and Fos-driven expression of tdTomato. **g**, Representative raster plots of recorded spike trains. Flash signs indicate periods of electric shocks. Red color indicates Fos-positive cells. **h**, Quantification of medium rank of mutual information with electric shocks from pooled neurons. Gray bars indicate shuffled null distribution. Rank-sum analysis, n = 21 fast-spiking neurons and 42 sparse-spiking neurons from 3 mice. **i**, Representative correlation matrix with recorded spikes. **j**, Quantification of mutual information to fast-spiking Fos-positive neurons, two-tailed t-test, n = 18 for Fos-positive connections and 33 for Fos-negative connections from 3 mice.

In order to directly test the above predictions, we performed *in vivo* voltage imaging in combination with electric shocks (**Figure 6e**). We were able to align field of view across days to track Fos expression (**Figure 6f**). Using a promoter that is active in both excitatory and inhibitory neurons, we observed clear populations with fast-spiking and sparse-spiking activities (**Figure 6g**), which were likely inhibitory and excitatory neurons, respectively. To examine the relationship between shock-responsiveness and Fos expression, we ranked the neurons by their mutual information between electric shock periods and recorded spike trains. We found that ranks of the Fos-positive fast-spiking neurons were significantly higher than the null distribution, indicating a strong correlation (**Figure 6h**). The sparse-spiking neurons, on the other hand, showed chance level ranking, consistent with calcium imaging data (**Figure 3**). To evaluate connectivity between neurons, we calculated mutual information between neuronal pairs using recorded spike data prior to the electric shocks (**Figure 6i**). We found that, on the population level, sparse-spiking Fos-positive neurons were weaker than Fos-negative neurons in connection to fast-spiking Fos-positive neurons (**Figure 6j**). Our *in vivo* voltage imaging data provided supportive evidence for the two predictions from our model. Together, these results indicate that interneurons may mediate the formation of the excitatory Fos-positive ensemble that is independent from their stimulus responsiveness.

## Discussion

Our study characterizes the relationship between information coding and Fos expression. Contrary to the previously established definition of engram cells^2^, we showed that excitatory Fos-ensemble does not represent coding of the experience during memory formation, but labels a distinct group of neurons for memory retrieval. In his 1904 book, German zoologist Richard Semon described a stimulus-responding brain process as memory engram, and considered the reactivation of memory engram as the basis of retrieval, which he termed ecphory^1,16^. Experimental evidence since about a decade ago has demonstrated that reactivation of Fos-expressing neurons induces memory retrieval, and named these neurons engram cells^4^. Our study provides unequivocal evidence that excitatory Fos-expressing neurons do not show experience-related activity during memory encoding and are recruited as a representation of the memory during retrieval. Considering previous studies showing that general suppression of DG activity during memory encoding blocked memory formation^18,19^, there is likely a separate neuron population in DG that are not Fos-positive and showed stimulus-specific activity patterns. Hence, our results suggest that memory ecphory is not the reactivation of stimulus-responding engrams, but rather these processes are supported by distinct neuron populations. And excitatory Fos-expression labels the specific population for ecphory.

We provide a potential mechanism based on inhibitory interneurons that assigns a specific memory to a Fos-expressing ecphory ensemble (analogous to the “binding” problem ^20^). While stimulus-triggered activities in excitatory neurons are too transient to elicit high intracellular calcium for Fos induction, fast-spiking interneurons may have a higher baseline calcium and thus are more likely to express Fos with stimulus. During post-event consolidation, these Fos-positive interneurons may drive selective inhibition, leaving the less-connected neurons more likely to express Fos. Therefore, the expression of Fos in excitatory neurons are not controlled by stimulus responsiveness, but are determined by their connectivity to interneurons^21^. This view is supported by our modeling as well as *in vivo* voltage imaging results, and is also consistent with previous work demonstrating that activities of interneurons during memory encoding could control the size of engram ensemble ^22^. Importantly, different types of interneurons with unique connectivity patterns may play different roles in this process and cell-type specific IEGs such as Npas4 ^23^(Neuronal PAS domain protein^24^) could confer more precise synaptic modifications that are necessary for the formation of excitatory Fos-positive ensemble. Alternatively, neurons in other brain regions that showed stereotypical responses to stimuli, could also be responsible for linking Fos-expressing ensembles to specific memories. Future studies that dissect this process are crucial for understanding the mechanisms of memory formation. Other possible mechanisms linking Fos expression and specific stimulus exist. Consistent with previous reports^25^, we found Fos-positive neurons generally showed elevated intracellular calcium. Although very high intracellular calcium activity could cause Fos expression, calcium activity triggered under physiological conditions had only a limited impact on Fos expression. In fact, we observed a small fraction of stimulus-coding neurons that also expressed Fos, although without preferrable probability. We did not rule out that this subpopulation is uniquely important in driving the Fos expression in the rest of the Fos-positive ensemble. For example, IEGs like Arc have been shown to spread mRNAs intercellularly via trans-synaptic connections^26^. While Fos induction via this type of mechanism has not been reported in the literature, our present study has not formally ruled out similar possibilities.

In summary, our research uncovers distinct neural processes underlying memory encoding and retrieval, and identifies excitatory Fos expression as a marker for memory ecphory. These findings clarify the interpretation of numerous previous studies using Fos-driven labeling, and raise previously unappreciated questions related to memory allocation and retrieval.

**Extended Data Fig.1:**
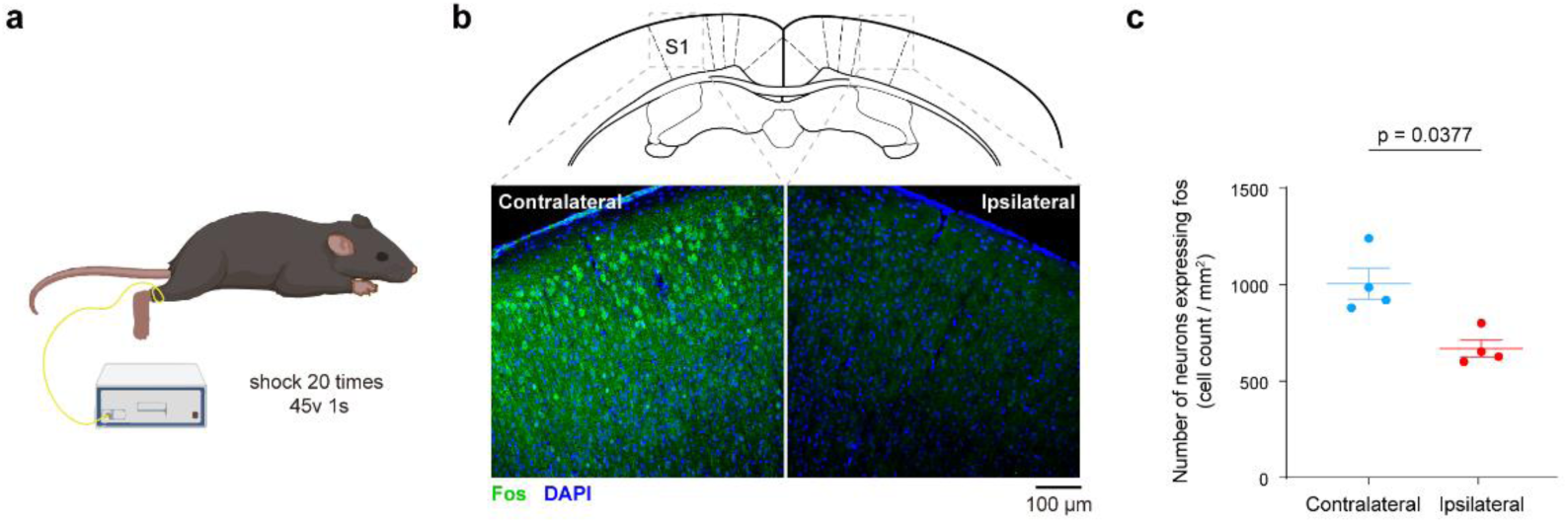
Unilateral electric shock in awake mice induced fos expression in the contralateral somatosensory cortex. **a**, Diagram of the electric shock procedure and conditions. **b**, Representative images of Fos immunohistochemistry from brains harvested 2 hours after electric shock. **c**, Quantification of fos expression on either side of brain slice. Data are presented as mean±S.E.M. Two-sided unpaired t test, n = 4 mice in each group.

**Extended Data Fig.2:**
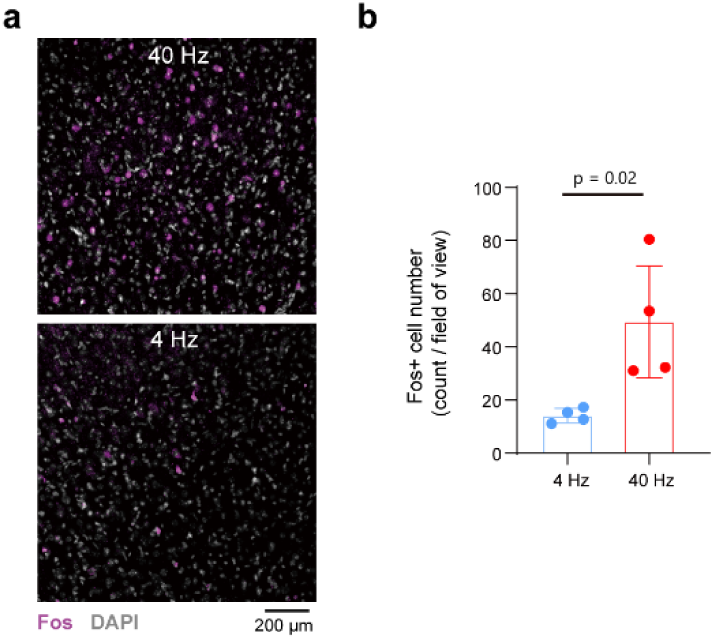
High frequency optogenetic stimulation induces fos expression. **a**, Representative confocal images of fos immunohistochemistry in cortical tissues from mice received 40Hz or 4Hz optogenetic activation. **b**, Quantification of fos-positive cell densities after 40Hz and 4Hz optogenetic activation. Two-sided unpaired t test, n = 4 mice.

**Extended Data Fig.3:**
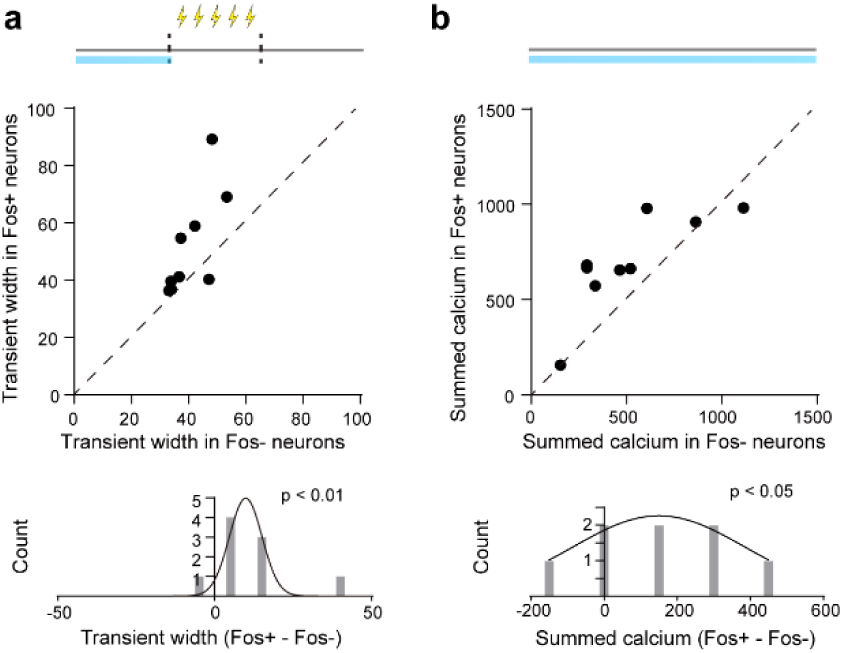
Fos-positive neurons show increased calcium activity prior to electric shock in somatosensory cortex. **a**, Calcium transient width of Fos-positive versus Fos-negative cells before the shock. Top, each circle indicates the mean across neurons for one mouse. The blue horizontal bar represents the time window for data analysis, covering the 10-minute period before electric shock. Bottom, session-wise difference histogram (grey) with kernel density estimation (black). Two-sided paired-sample t test, n = 9 mice. **b**, Calcium transient width of Fos-positive versus Fos-negative cells without shock. Top, each circle indicates the mean across neurons for one mouse. Bottom, session-wise difference histogram (grey) with kernel density estimation (black). Two-sided paired-sample t test, n = 8 mice.

**Extended Data Fig.4:**
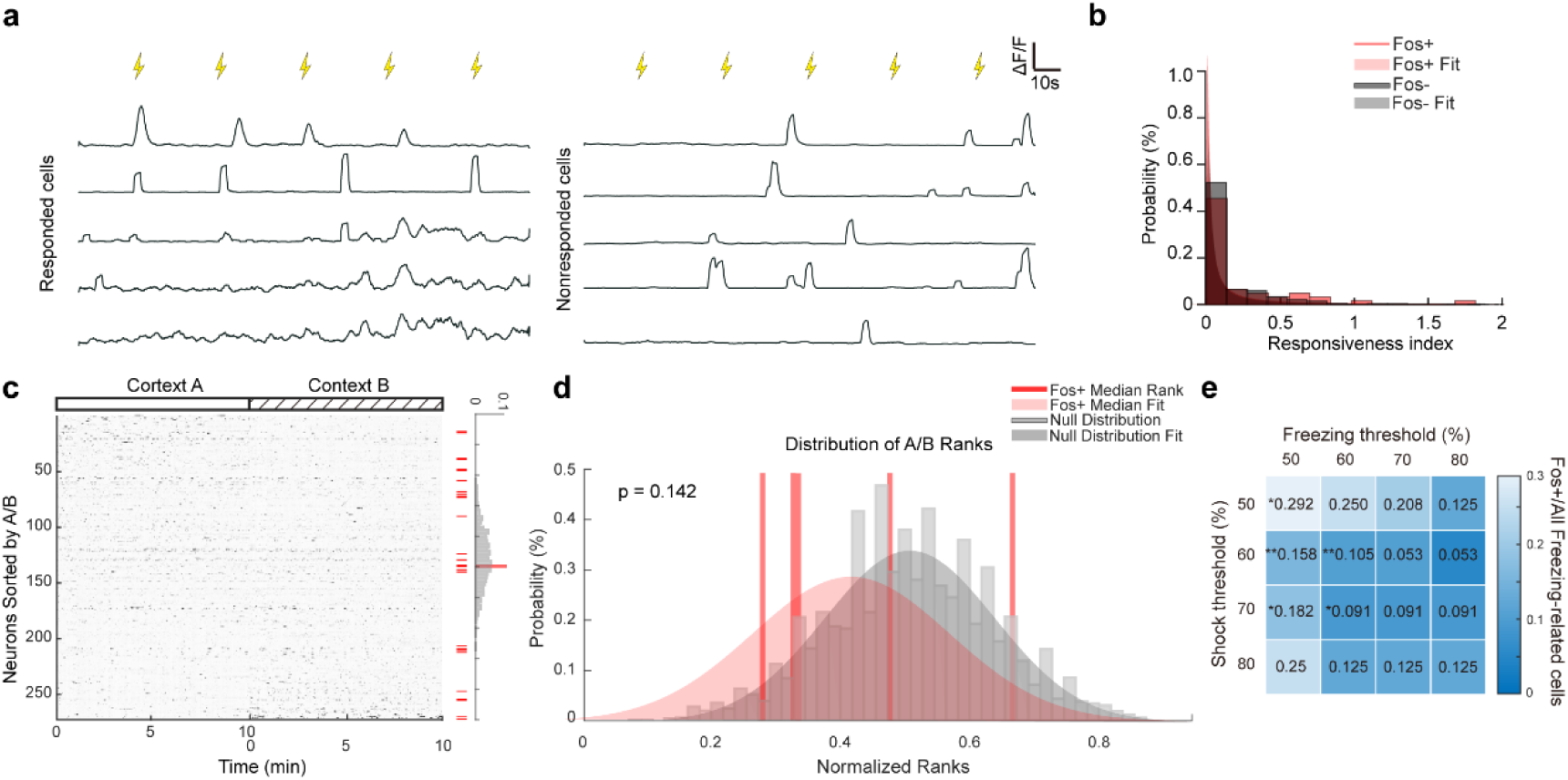
Activity of Fos-positive cells in DG does not encode stimulus. **a**, Representative calcium traces from shock responsive and non-responsive cells in DG. Yellow flash signs indicate the time of electric shock. **b**, Histogram of the responsiveness index in fos-positive and fos-negative neurons. Shades indicate fitting results of log-normal distributions. **c**, Example of neuronal activities from the adaptation day imaging session, ranked by the mean activity ratio between A and B contexts, with fos-positive neurons labeled with red bars. Right panel shows the medium rank from fos-positive neurons, and null distribution generated by random sampling the same number of neurons. **d**, Quantification of medium rank from individual mice, with shades indicate fitting results of normal distribution. Rank-sum analysis, n = 5 mice. **e**, Proportion of Fos-positive cells among cells responsive to shock and freezing at various percentiles threshold of the mutual information. Double-responsive cells show disproportionally low probability for fos expression. Chi-square tests for each threshold condition.

**Extended Data Fig.5:**
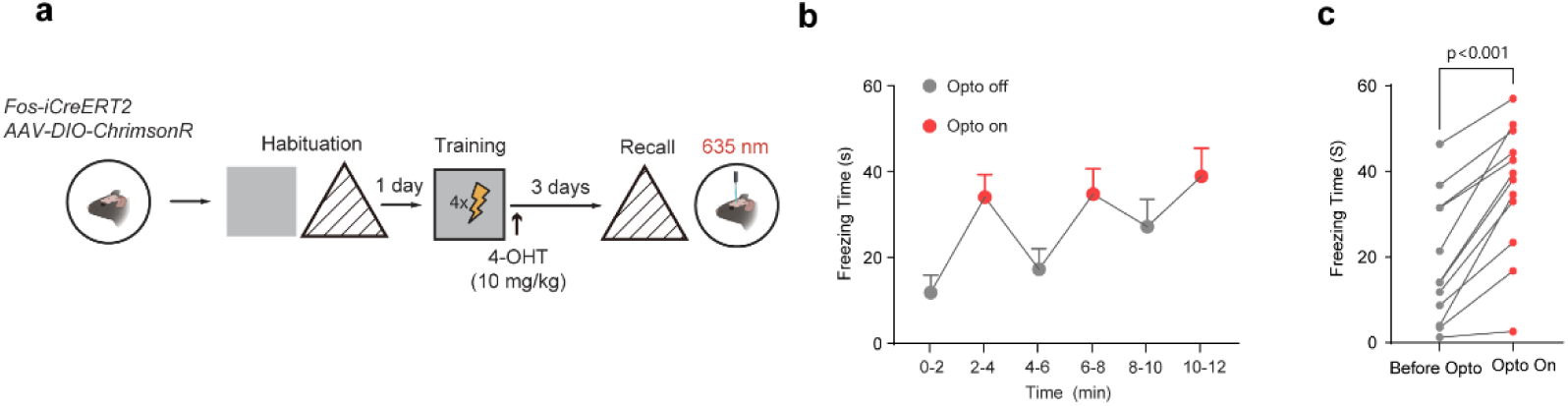
Optogenetic activation of fos-positive cells induces memory-guided freezing behavior. **a**, Schematics of the experimental design and timelines. **b**, Quantification of the freezing behaviors with (red shades) and without optogenetic activation. Data are represented as mean±S.E.M. **c**, Quantification of the freezing time in the test session with and without optogenetic activation. Paired t test, n = 12 mice.

**Extended Data Fig.6:**
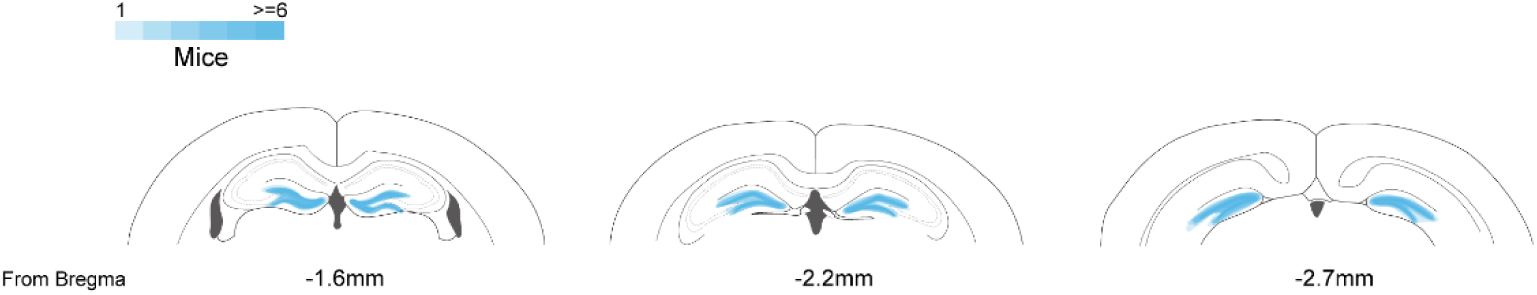
Virus expression map in the hippocampal DG region. **a**, Diagram of the viral infection regions at different anterior-posterior positions. Shades indicate the spread of virus as assessed by fluorescence.

## Material and methods

### Mice

Trap2 (IMSR_JAX:030323, The Jackson Laboratory) mice were used in this study. Ai9(IMSR_JAX:007909, The Jackson Laboratory) mice were crossbred with Trap2 mice. The genotyping of Trap2 mice was performed according to the instructions provided by The Jackson Laboratory. All of the animal procedures were approved by the Institutional Animal Care and Use Committee at Fudan University.

### Virus injection

Before injecting virus, mice were anaesthetized with isoflurane and a craniotomy was made around 1.5 mm lateral to the midline (right hemisphere) and 0.58 mm posterior to bregma. Virus injections were performed using beveled glass micropipettes ∼0.5 mm below the dura, connected to an oil-filled pressurized injector (Nanoinject III, Drummond Scientific) at 1 nL/s. When the injection finished, pipettes were kept in the injection site for 5 minutes before retraction. For population imaging in Trap2 mice, 250 nl of AAV2/9-CaMKII-GCamP6f (3 × 10^12^ genome copies per ml) was injected at each site. For imaging voltage signals and calcium activities simultaneously, 400nl of AAV2/9-CaMKII-jRGECO1a (3 × 10^12^ genome copies per ml) and 400nl of AAV2/9-CaMKII-ASAP4 (1 × 10^13^ genome copies per ml) were injected sequentially at each site. For optogenetics infection, 300 nl of a 1:1 mixture of AAV2/9-CaMKII-GCamP6f (3 × 10^12^ genome copies per ml) and AAV2/9-hsyn-ChrimsonR (4.93 × 10^12^ genome copies per ml) was injected into each location.

### Cranial window implant

Trap2::Ai9 mice were anaesthetized with isoflurane and hair was removed from the skull area. Dexamethasone (2 mg per kg) and carprofen (5 mg per kg) were given subcutaneously at this point. The mouse was placed on a heating pad during the surgery and anesthesia was checked periodically. Povidone-iodine solution was applied to the skin and cleaned with ethanol, and eye ointment was applied to the eyes. A small piece of skin was removed to expose the skull, and the membranous layer on the skull surface was removed by forceps. A 3 mm diameter circle was drilled on the hemisphere of virus infusion (approximate location of the centre is −0.58 mm from bregma and 1.5 mm from the midline). The skull was rinsed with sterile PBS periodically to avoid excessive heating. The skull was thinned in a circumferential area and then lifted with fine forceps without causing injury to the underlying pila surface. Gelfoam sponge was used to absorb blood after lifting the skull. Using a pair of very fine forceps, the dura was removed within the circle area and a 3 mm cover glass was gently pressed onto the brain surface and glued to the skull. A customized head bar was chronically implanted onto the skull. For chronic imaging, mice were placed onto a heating pad to recover after the surgery and given dexamethasone (2 mg per kg) and carprofen (5 mg per kg) for 3 days. Imaging procedures started 1 month after the surgery.

### In vivo two-photon imaging

In vivo imaging was performed using a two-photon microscope (Thorlabs, Bergmo II). Two-photon excitation was achieved by a Ti:sapphire femto-second laser (Coherent, Chameleon Vision II). GCaMP6f was excited at 920 nm; tdTomato were excited at 1,000 nm; jRGECO1a and ASAP4 were co-excited at 985nm. The excitation laser power was below 25 mW. Imaging was carried out with a water immersion objective (Nikon, 0.8 numerical aperture, magnification 16x). We used a custom-designed headplate holder designed for the repetitive daily mounting of mice on the treadmill. For chronic imaging, a location close to the center of the cranial window was selected as the starting point and the blood vessel pattern was recorded. The coordinates of each ROI were recorded as well. To relocate in the next imaging session, the starting point was relocated on the basis of the recorded coordinates and the field of view was adjusted to match the recorded blood vessel pattern. In each imaging session, calcium activities of neurons were recorded at 15 Hz for 30 min. Simultaneous imaging of voltage signals and calcium activities were recorded at 116 Hz.

### Calcium trace analysis

Raw image files were processed with a motion correction pipeline (NoRMCorre: An online algorithm for piecewise rigid motion correction of calcium imaging data) and cellular signals were analyzed on a 2x spatially downsampled and 3x temporally downsampled data with a previously established algorithm (Fast and statistically robust cell extraction from large-scale neural calcium imaging datasets). Extracted cell signals were manually inspected to eliminate artifacts. To analyze calcium transients’ properties, we applied a custom-made script to isolate individual transient using a z-score based threshold method. The peak amplitude and width of each transient were derived and averaged across all transients from each cell. To align data cross sessions, we manually set control points of unique blood vessel patterns in the field of view as anchors for alignment, and performed linear transformation of images. The cell masks from the first session were transformed onto the images of the second session, and the fluorescent intensity of the red channel were measured as indication of Fos expression. To train and test classifier models using extracted calcium transient features for predicting Fos expression, we selected random and balanced subset of cells from Fos-high and Fos-low groups in each mouse. Among the selected cells, 80% were used as training set for a support vector machine (SVM) model with polynomial kernel function, and the remaining 20% were used as testing set to evaluate the performance of the model. To evaluate the responsiveness to electric shocks, we compared the calcium activity 10-frame before and after electric shocks and calculated the statistical significance for each cell (p1). In addition, we calculated the mutual information between calcium traces with electric shock periods and compared to a null-distribution generated from shuffled data (p2). A rank-based statistical analysis was applied with both p1 and p2, with false discovery rate 0.05 as correction for multiple comparison. For analysis of noise correlation, the max amplitude of each electric shock period for each cell was calculated, and we further computed the differences of each shock to the mean amplitude. Pearson’s correlation coefficients of these differences were calculated between each pair of cells. The median number from all coefficients was used as the representative noise correlation from each mouse.

### Dimension reduction for calcium transients and neuronal activities

To visualize the population-level difference of calcium activates between Fos-high and Fos-low neurons, we reduce the four calcium transient features for each neuron into two-dimensional presentation by Uniform Manifold Approximation and Projection (UMAP). For each mouse, we calculated the centroid coordinates of Fos-high and Fos-low neurons. The vector pointing from the Fos-low centroid to Fos-high centroid was projected onto the primary component of the UMAP coordinates. The predicted probability of Fos expression for each cell was estimated by normalizing the score from a SVM model (see **Calcium trace analysis**). To validate the Fos-expression estimation, the sum of predicted probability and count of Fos-high neurons were assessed within a sliding window (side length = 0.5 or 1.5). The accuracy was determined by measuring the linearity between summed probability and the Fos-high neuron counts. To visualize neuronal population activity, we reduced the N-dimensional activities to two dimensions using Partial Least Squares (PLS), which was supervised by prior-freezing state.

### Template construction for population-level neuronal activity

We constructed time-wise averaged templates representing neuronal activity during prior-freezing period and the initiation of freezing behavior. The prior-freezing template was an N-dimensional vector generated by averaging the neuronal activities over a one-second period preceding each freezing trial. The freezing-template was created by averaging the neuronal activity during the initiation phase of freezing behavior. To quantify the representation of the initiation phase, we defined freezing-average and resting-average template as the averaged neuronal activities during behaviorally freezing and resting period, respectively. The state representation at each time point was characterized by the Mahalanobis distances between neuronal activity and these two templates. Specifically, the freezing behavior initiation phase was identified as a rapid shift in state representation from resting to freezing.

### Rank distribution test of Fos expressing neurons

During the shock phase, an SVM model with linear kernel was trained using neuronal activities as input and shock stimulation as labels. The SVM coefficient assigned to each neuron served as the measure of the neuron’s relevance to the stimulation. Neurons were ranked by their SVM coefficients to establish shock ranks. During the retrieval phase, neurons were ranked according to their orders in the freezing or prior-freezing template (see **Template construction for population-level neuronal activity**). To test whether Fos-high neurons showed greater relevance in shock phase or retrieval phase, Wilcoxon rank sum test was applied to the ranks of Fos-high neurons and a rank distribution under the null hypothesis generated by random sampling.

### Hind limb electric shock

To apply electric shock, we attached copper wires to one of the hind limbs in the head-fixed mouse. Another stream of wires was connected to the head bar on the same body side of the mouse. To avoid short-circuiting, the imaging setup was insulated from the air table by plastic connectors. Electrical stimulation was provided by a direct current shocker (ISO-Flex, A.M.P.I.) at the output voltage of 40 volts. Output voltage was verified before the experiment on each session. At this strength, mice did not show violent movement towards the shock. We applied 5 shocks of 1-second duration, with a 30-second interval between shocks. Electric shock was accompanied by a TTL signal that was sent to an Arduino board to synchronize with imaging.

### Optogenetics

Optogenetic laser was provided with a desktop source (Thinker Tech, QAXK-LASER-R1) with a fixed 635 nm laser. Laser was directed to the cranial window with multi-mode optic fibers while the mouse was head fixed. For optogenetic stimulation, the output of the laser was measured and adjusted to 5 mW before each experiment. The pulse onset, duration, and frequency of light stimulation were controlled by a programmable pulse generator attached to the laser system. Optogenetic stimulation was usually performed 3 weeks after AAV injection. In order to minimize variability between individual mice, we used a design of stimulating the same mouse’s two hemispheres with different frequencies, so that we can compare the effect within the same mouse. In addition, to avoid the effect of inter-hemispherical projections, we chose S1 and M1 on different hemispheres, and randomly assigned the brain regions with different stimulation protocols. Two stimulation protocols were tested after validation from brain slice electrophysiology and *in vivo* imaging: in every 30 seconds, we applied a 10-second stimulation of 40 Hz or 4 Hz light pulses followed by a 20-second period without light. Each light pulse was set at 12.5 ms duration. The 40 Hz group received a total of 2-minute stimulation and the 4 Hz group received 20 minutes so that the total number of light pulses were kept the same between groups.

### Immunohistochemistry

Mice were anesthetized with isoflurane and sequentially perfused with saline and PBS with 4% PFA. Whole brain tissue was cut into coronal sections of 45-micron thickness using a vibrating microtome (Leica, Germany), incubated overnight with blocking solution (PBS containing 5% donkey serum and 0.2% Triton X-100), then treated with the anti-fos primary antibody (Abcam, ab190289) diluted with blocking solution for 2 h at room temperature. Primary antibodies were washed five times with washing buffer before incubation with secondary antibodies (Abcam, ab150073). Sections were counter stained with DAPI and mounted on cover slides. Sections were imaged with a Nikon confocal microscope (10× objective lens). Samples were excited by 405- and 488-lasers in sequential acquisition mode to avoid signal blead through. Saturation was avoided by monitoring pixel intensity under Hi-Lo mode. Confocal images were analyzed with ImageJ software.

### Slice preparation and electrophysiological recording

C57BL/6J mice were sacrificed by rapid decapitation and brain tissues were immediately dissected out. In ice-cold sucrose-based slicing solution (normal ACSF listed below but with NaCl replaced with equiosmolar sucrose) that had been bubbled with 95% O2 and 5% CO2, tissue blocks from mice were cut coronally with a vibratome (Leica VT1000S). Cortical slices (300-μm thick) were collected and incubated at 35°C in aerated artificial cerebrospinal fluid (ACSF) containing (in mM): NaCl 126, KCl 2.5, MgSO4 2, CaCl2 2, NaHCO3 26, NaH2PO4 1.25, and dextrose 25 (315 mOsm, pH 7.4). After 90-min incubation, slices were then incubated at room temperature until use.

Slices were transferred to the recording chamber and perfused with aerated ACSF (34-35°C) at a rate of 1.2 ml/min. Cortical neurons were visualized under upright infrared differential interference contrast microscope (BX51WI, Olympus). The impedances of patch pipettes for somatic recordings were 5–7 MΩ, when filled with the internal solution (in mM): K-gluconate 145, MgCl2 2, Na2ATP 2, HEPES 10, EGTA 0.2 (286 mOsm, pH 7.2). We selected cells that expressed ChrimsonR with red fluorescence. ChrimsonR was excited using the same laser source and energy that we used for *in vivo* experiment, and the CED Power 1401 system was employed to set the signal of the light source, marked on the whole-cell patch-clamp recording. The light was administered for 5 milliseconds as the duration for each stimulation.

### Miniscope imaging

Our virus injection and lens implantation were performed concurrently on the same day to minimize the potential misalignment between the lens and virus injection sites, which can occur after prolonged recovery. This approach prevents imaging failures related to the misalignment of the focal plane. The procedure followed our previously published protocol ^14^. Briefly, mice were pre-treated with carprofen (5 mg/kg, subcutaneous injection) prior to surgery to mitigate inflammation and brain swelling. A craniotomy was performed over the right dental gyrus (DG, AP: -2.18, ML: +1.5), followed by careful removal of dura. Cortical tissue was gently aspirated with continuously application of cold Ringer’s solution. A 1-mm diameter gradient refractive index (GRIN) lens (CLHS100GFT003, GoFoton) was implanted at a depth of -1.7 mm from the skull surface and secured to the skull using light-curable glue. A custom-made metal head bar was subsequently attached for head fixation. The baseplate was installed two weeks post-surgery. Mice were allowed to recover for 4-6 weeks before imaging experiments. Imaging was carried out with a dual color miniscope system (Inscopix, nVue). Blue excitation light was set at approximately 0.2 mW, and the detector was configured with a gain of 1.8, and a sampling rate of 20 Hz. During the recall phase, a single snapshot was taken to collect red fluorescence immediately prior to imaging in order to capture the *Fos*-driven expression of tdTomato.

### Fear conditioning

The fear conditioning experiment is a three-day setting. On the first day, mice were placed in Context A (a small chamber with metal bar flooring, 31×24×21 cm; MED Associates) and Context B (a smooth plastic floor with a striped background) for habituation, with a minimum 5-hour interval between the two sessions. On the second day, during the training phase, the mice were placed in the fear-conditioning chamber (Context A) and subjected to five foot shocks (0.7 mA, 1-second duration, 1-min interval). Twenty-four hours after training, the mice were returned to Context A for a 10-minute contextual memory test by assessing freezing behavior. This was followed by a 10-minute exposure to Context B for evaluating their freezing behavior in a non-threatening context.

For experiments involving optogenetic inhibition of fos-expressing cells, mice were divided into three groups:

First group: Control group. On day one, the mice were placed in an irrelevant environment (Context C). Optogenetic activation (30 seconds) was used to stimulate neurons sparsely labeled in the hippocampal DG region, marking high-fos cells in an unrelated environment.

Second group: fos-assigned group. On day two, five minutes prior to exposure to Context A (unrelated environment), 30 seconds of optogenetic activation was used to stimulate high-fos cells in the hippocampal DG region, artificially associating these cells to Context A.

Third group: Shock-inhibition group. On day two, 30 seconds of optogenetic activation was performed before the mice were placed in Context A to assign high-fos cells to this context. During the foot shocks, optogenetic inhibition (470 nm, blue light, 5 mW, 7 seconds) to inactivate the high-fos cells during the shock.

### Freezing behavior detection

Noldus EthoVision XT was used to analyze freezing behavior in mice. Videos were calibrated to the actual chamber area, and background subtraction was applied for subject detection. The zones of interest were defined, and detection parameters were adjusted for optimal tracking. Freezing was defined as a lack of movement below 0.5 cm/s for at least half second. Analysis provided data on freezing state of each video frame. Data were exported for further statistical evaluation. Behavior detection were verified against video playback to ensure accurate identification of freezing events.

### Leak-integrate-fire model

The microcircuit model was implemented using MATLAB (2021b). Neuronal membrane potentials were calculated using the “Leak-integrate-and-fire” (LIF) model, described by the following differential equation:

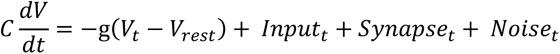

Vt denotes the membrane potential at timepoint t, Vrest is the resting membrane potential set at –60 mV, and Inputt, Synapset, and Noiset represent the input signal, summed synaptic inputs, and noise (modeled using a Poisson distribution), respectively. The parameters C and g denote the cell’s capacitance and conductance, with C/g representing the time constant of membrane potential changes, set to 10. For excitatory neurons, firing threshold is set at -45 mV, followed by a reset to - 70 mV. For inhibitory neurons, firing threshold is set at -55 mV, followed by a reset to -65 mV. The model simulated 100 neurons, including 30 interneurons and 70 excitatory neurons. We assumed that interneurons form synaptic connections with the excitatory population at a probability of 0.3. Excitatory neurons form connection among themselves at a probability of 0.1. Synaptic inputs were calculated by summing synaptic currents from connected neurons activated in the preceding timepoint, with excitatory weights initialized at 1.25 and inhibitory weights at –0.45. To simulate electric stimulation, we randomly selected 20 cells and added an input of 10 mV to these cells for 10 timepoints for 5 times. After generating the simulated spike trains, we then modeled the intracellular calcium using similar differential equation:

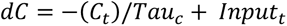

Where the time constant for calcium decay (Tauc) is set at 50. Inputt indicates the firing status of the neuron with a multiplier coefficient of 10. Interneurons with calcium concentration above 150 or excitatory neurons above 120 will be designated to express Fos.

Fos expression will lead to an increase in the excitability as modeled by a decrease of firing threshold by 2 mV. In addition, the synaptic connections from Fos-positive neurons will have a 5-five amplification of their weights. The offline broad activation is modeled as an input current of 6 mV to 80 random cells that lasts for 100 timepoints. After generating the spike trains, we then calculated the intracellular calcium following the same procedure and determined the Fos expressing cells again. We ran the models for 100 times, and the average results were recorded.

## Data availability statement

All raw data generated for this study are available from the corresponding author upon request.

## Code availability statement

All scripts used in this study are available on Github: https://github.com/PaulYJ/FOS-ecphory, and are free to use and share freely.

## Author contribution statement

P.Y. and N.D. designed the study. N.D., CY.W., YT. L. performed the in vivo imaging and optogenetics experiments. Y.X. and Y.S. designed and performed the electrophysiology experiments. P.Y., Q.S. and N.D. analyzed the data. X.W. maintained the transgenic mice used in the study and helped with the illustration. P.Y. and N.D. wrote and edited the manuscript.

## Acknowledgement

We express our sincere gratitude towards Dr. Yingxi Lin (UT southwestern) for her valuable feedback of our study. This study was supported by Shanghai Pilot Program for Basic Research – FuDan University 21TQ1400100 (22TQ019), Shanghai Municipal Science and Technology Major Project, the Lingang Laboratory (grant no. LG-QS-202203-09) and National Natural Science Foundation of China (32371036). Y.X. was additionally supported by National Natural Science Foundation of China (32200951) and China Postdoctoral Science Foundation (2022M720801). Y.S. was additionally supported by National Natural Science Foundation of China (32130044 and T2241002), and STI2030-Major Projects (2021ZD0202500).

